# Visualizing Amino Acid Substitutions in a Physicochemical Vector Space

**DOI:** 10.1101/2021.07.15.452549

**Authors:** Louis R. Nemzer

## Abstract

A three-dimensional representation of the twenty proteinogenic amino acids in a physicochemical space is presented. Vectors corresponding to amino acid substitutions are classified based on whether they are accessible via a single-nucleotide mutation. It is shown that the standard genetic code establishes a “choice architecture” that permits nearly independent tuning of the properties related with size and those related with hydrophobicity. This work sheds light on the non-arbitrary benefits of evolvability that may have shaped the development standard genetic code to increase the probability that adaptive point mutations will be generated. Illustrations of the usefulness of visualizing amino acid substitutions in a 3D physicochemical space are shown using recent datasets collected regarding the SARS-CoV-2 receptor binding domain. First, the substitutions most responsible for antibody escape are almost always inaccessible via single nucleotide mutation, and change multiple properties concurrently. Second, it is shown that assays of ACE2 binding by sarbecovirus variants, including the viruses responsible for SARS and COVID-19, are more easily understood when plotted with this method. The results of this research can extend our understanding of certain hereditary disorders caused by point mutations, as well as guide the development of rational protein and vaccine design.

## Introduction

The canonical genetic code, which maps three-letter DNA codons into amino acids, has long been recognized to possess intrinsic error-correcting mechanisms. In particular, the encoding is nearly optimal^1^ in minimizing the chance that a single nucleotide mutation will cause a drastic change in the physicochemical properties of the resulting residue.^2^ One celebrated example is that codons with the middle nucleotide U almost always produce a hydrophobic amino acid. However, the specific constraints of allowed substitutions, which lead to new variants, have received comparatively less attention. There is evidence that the genetic code has been shaped by objectives besides robustness to error.^3,4^ In addition to the adaptive value in avoiding nonconservative mutations, natural selection may favor a system in which the most important^5^ physical and chemical properties of proteins can be adjusted independently. One reason this topic has been less studied is that it raises philosophical meta-questions surrounding how to think about the selection pressure on systems to exhibit “evolvability,”^6^ which is the ability to adapt efficiently to changing environments.^7^ It should be remembered, however, that the standard codon table is itself a product of evolution,^8,9^ albeit over a much longer timescale^10^ compared with the evolution of individual genes. Selection pressures on this “choice architecture”^11,12^- the system of presented options - would favor systems with the capacity to generate novel heritable variation^13,14^ by appropriately constraining the impact of any single point mutation. That is, potentially valuable changes should generally significantly alter at most one physicochemical property. Conversely, pathogens attempting to disguise themselves to evade the immune system of the host would benefit from amino acid substitutions in antigens that greatly change their binding with antibodies. This work is among the first assess the canonical genetic code in this light.

## Background

The physical and chemical properties of the twenty standard amino acids provide the raw materials for all biological protein structure. Several hereditary disorders are linked to specific single amino acid substitutions that break or substantially change the function of the resulting protein. Recent advances in genetic editing, such as CRISPR, have opened the possibility for targeted therapeutic interventions. Additionally, the products of rational protein design, including the SARS-CoV-2 prefusion stabilized spike protein^15^ are already in clinical use, but this method depends on detailed knowledge the effects of amino acid substitutions.

Complex adaptive systems, from organisms to societies, require “evolvability,” which is an ability to produce new traits,^16^ but almost always^17^ via bounded tinkering.^18, 19^ The guiderails that constrain these changes are generally stable over very long periods compared even with evolutionary time. Using an analogy from political systems, a written Constitution provides the metarules by which regular statutes may be altered. This Constitution may itself be modified with via amendments, but much more slowly and deliberately. Similarly, the canonical codon table and fixed amino acid properties provide the context for the subset of mutations that are accessible by a single-nucleotide DNA mutation, which can be considered essentially stationary even over evolutionary timescales.^20^ The near universality of the current genetic code evinces its slow rate of changes.^21^

The sum total of these constraints comprises a choice architecture. In human-designed systems, a choice architecture can include everything from limits on the legislative branch, mandatory failsafe interlocks, or simply requiring extra clicks to get a custom software installation.^22^ From a genetic perspective, there is an adaptive advantage to using a codon code in which the most important physical and chemical properties can be independently adjusted. Protein evolvability may be quantified in various ways, such as the proportion of sites shown to be under positive selection multiplied by the average rate of adaptive evolution at these sites.^23^

While error-resistance is clearly an important feature, since beneficial variants are likely to be similar to existing proteins,^24^ a purely static genome would not be conducive to long term evolutionary success in constantly shifting environments.^25^ Some previous work on protein evolvability under the genetic code^26^ examined the efficiency of allowed mutations as a “search algorithm”^27^ that explores a larger fraction of the space of functional variants when compared with random codes.^28^ It has also been shown that the substitutions accessible via point mutation in the TEM-1 gene for beta-lactamase were more likely to be adaptive than inaccessible changes.^29^ Also, the stepwise evolvability of proteins depends on the connectivity^30^ of amino acid sequence space.^31^ Some previous research considered decision trees^32^ or random forest algorithms^33^ for point mutations using physicochemical distance as part of the prediction^34^ of deleterious effects. However, this research is one of the only projects to consider the directionality of the vectors, since edges oriented (nearly) parallel to one of the axes would primarily change only one physicochemical aspect at a time. As an example of the power of this framework, recent data on the amino acid substitutions in the SARS-CoV-2 receptor binding domain associated with immune escape are analyzed. In addition, when visualized in a similar space, the results for site-saturation mutagenesis ACE2 binding assays with sarbecoviruses can be interpreted as either idiosyncratic – such as significantly reduced binding when a charged amino acid is introduced – or more general, as when based on size and hydrophobicity.

## Method

This work builds on previous results, including a principal component analysis (PCA) used to generate a 3D representation in physicochemical space (figure 1),^35^ as well as the classification system for DNA mutations.^36^ Here, only nonsynonymous missense mutations are considered, and when combining forward and backward changes together, there are (20 x 19)/2 = 190 possible amino acid substitution pairs. Of these, 75 are accessible via point mutations in the canonical genetic code, and 115 are not. This leads to a vector space (see Table 1) in which the x-, y-, and z-axes are assigned to size, hydrophobicity, and charge, respectively. The undirected edges represent substitutions separated into “possible” and “not possible,” as shown in figure 2. The thickness each edge in the “possible” network represents the number of permutations that can give rise to that substitution (Table 2). The coloring of the edges shows how damaging the change is predicted to be, on average, based on the BLOSUM62 substitution matrix.^37^ To further motivate the usefulness of this vector space representation, some notable single-nucleotide mutations are shown in figure 3. These include those in β-globin gene causing sickle cell anemia, the HFE gene causing hereditary hemochromatosis, the altered EphB2 gene in all crested pigeons,^38^ and a SARS-CoV-2 variant noted for its ability to evade the immune system. The possible substitutions are then separated by the mutated nucleotide and mutation type (figure 4). The insets provide the legend for the mutation types, based on the property shared by the original and new nucleotide (puRine, pYrimidine, Weak, Strong, aMino, or Keto). The order is based on the importance of the codon position,^39^ in which the middle nucleotide is the most determinative, followed by the first and then the third. The joint histograms of the BLOSUM62 and Euclidian PCA distance is separated into possible and not possible (figure 5).

**Figure 1:**
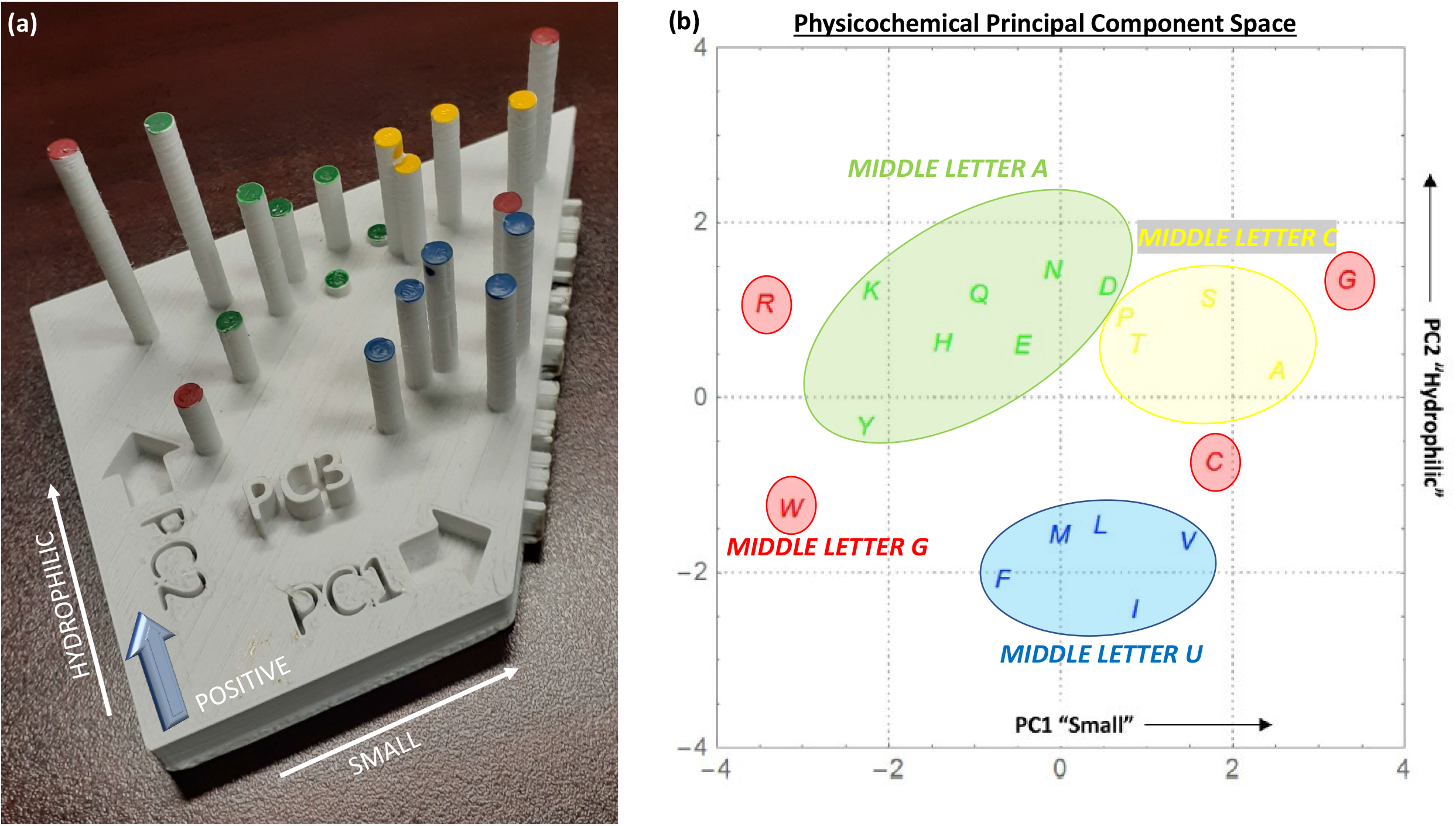
(**a**) 3D-Printed visualization of the physicochemical vector space generated by principal component analysis (PCA). The first principal component (PC1) is identified with the size of each amino acid, with smaller values to the right. PC2 is assigned to the hydrophobicity, with more hydrophilic residues at the top. The vertical height of each bar represents PC3, which shows the charge the amino acid, with more positive values being taller. (**b**) Two-dimensional projection of the same data. In both panels, the color indicates the middle nucleotide of the codon that encodes for each amino acid.

**Figure 2:**
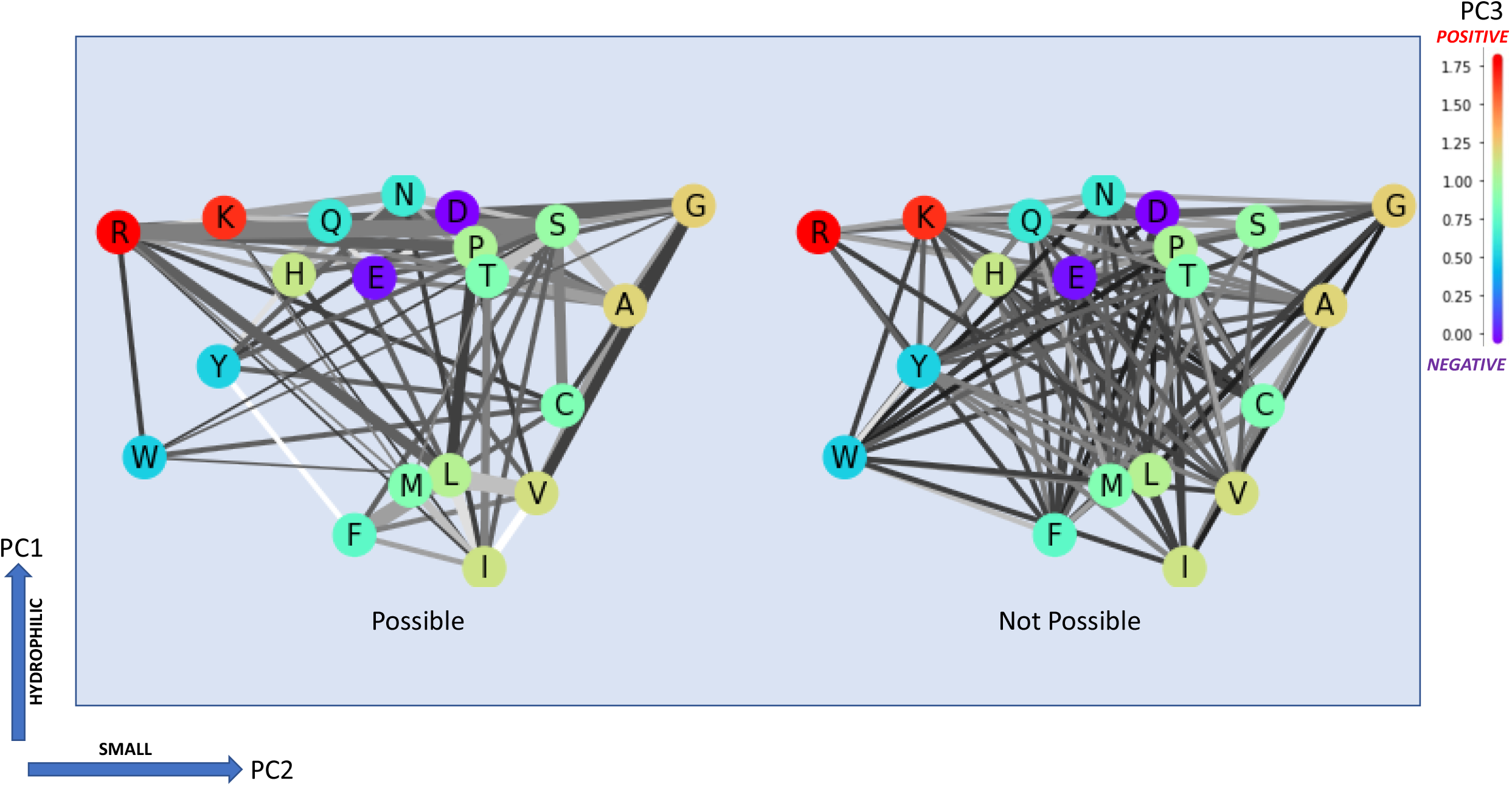
PCA diagrams in which the undirected edges display all possible amino acid substitutions. PC1 and PC2 are represented by the x- and y-axes, respectively. The PC3 value is indicated by the color of the node, shifted so the minimum is zero (see colorbar). The left figure shows substitutions that are accessible via single-nucleotide (point) mutations, with the thickness showing the number of possible codons associated with it. Conversely, the figure on the right has the remaining inaccessible substitutions that require more than one nucleotide change. The colors of the edges represent the impact of the substitution based on the BLOSUM62 matrix, with darker lines being more potentially harmful.

**Figure 3:**
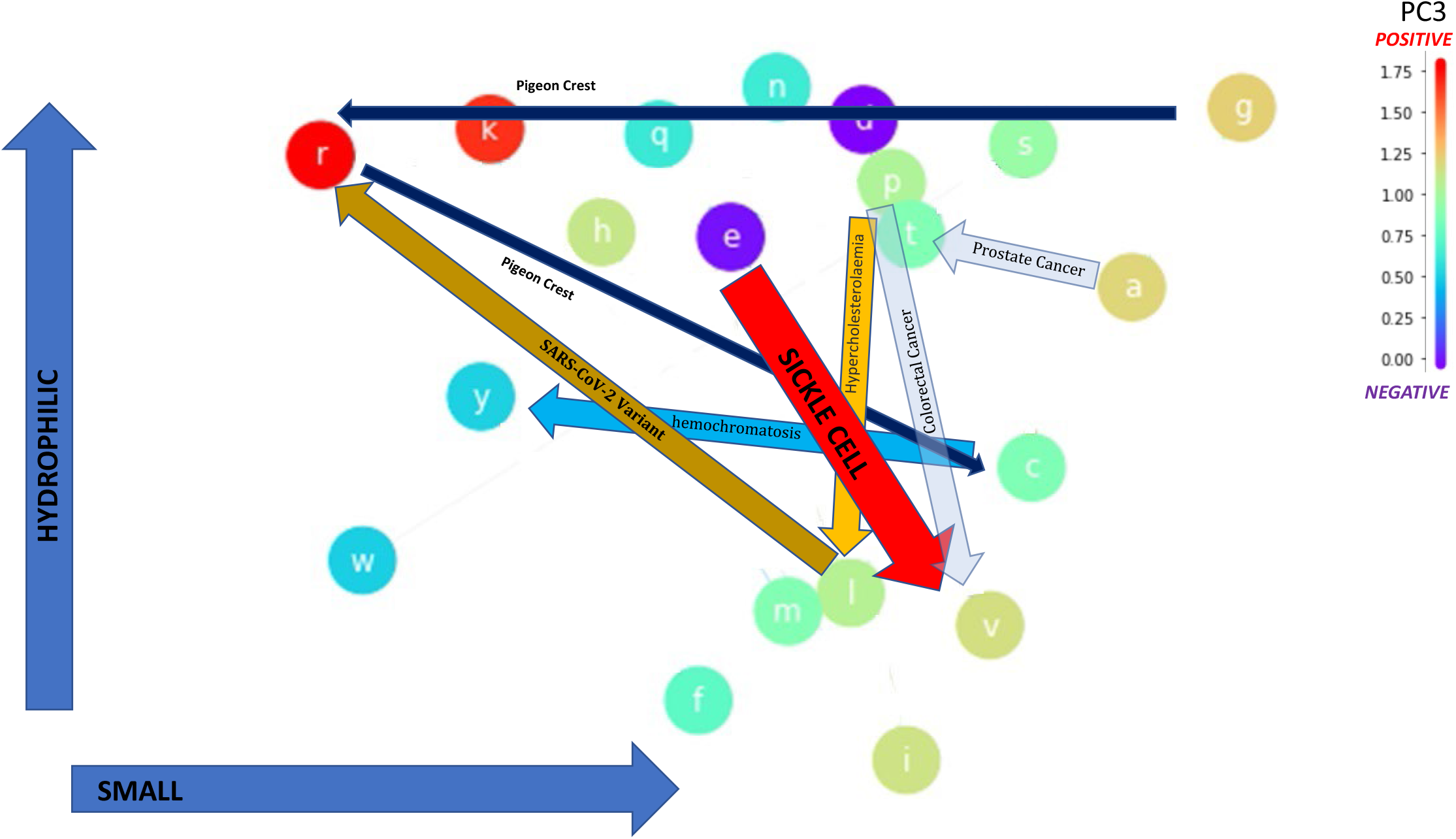
Selected notable point mutations. In some cases, a single missense mutation destroys the function of the resulting protein (e.g., in the kinase domain of EphB2 in crested pigeon), while in others the behavior is significantly altered (e.g., sickle-cell hemoglobin). One SARS-CoV-2 variant shown to have high immune escape^57^ is also included.

**Figure 4:**
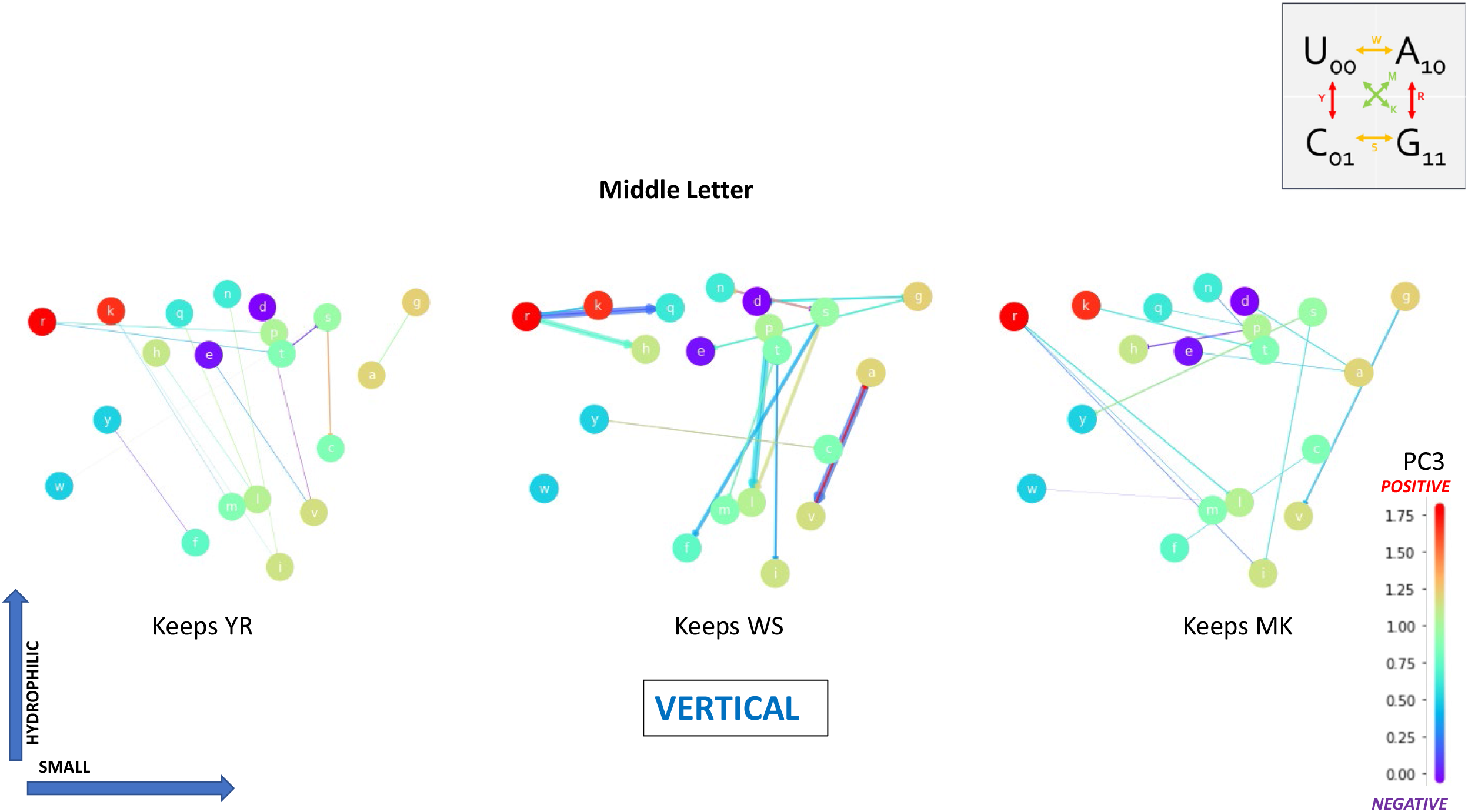
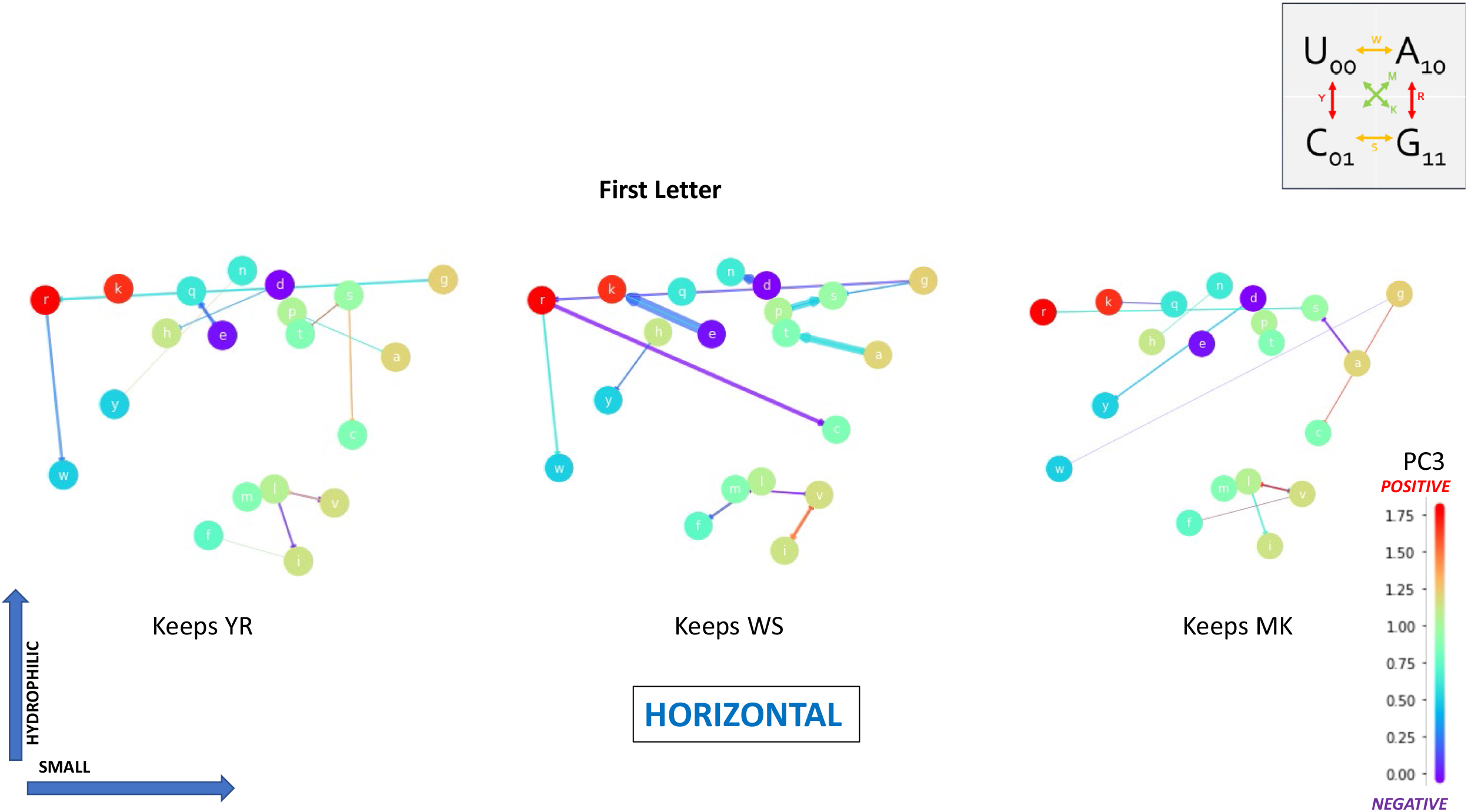
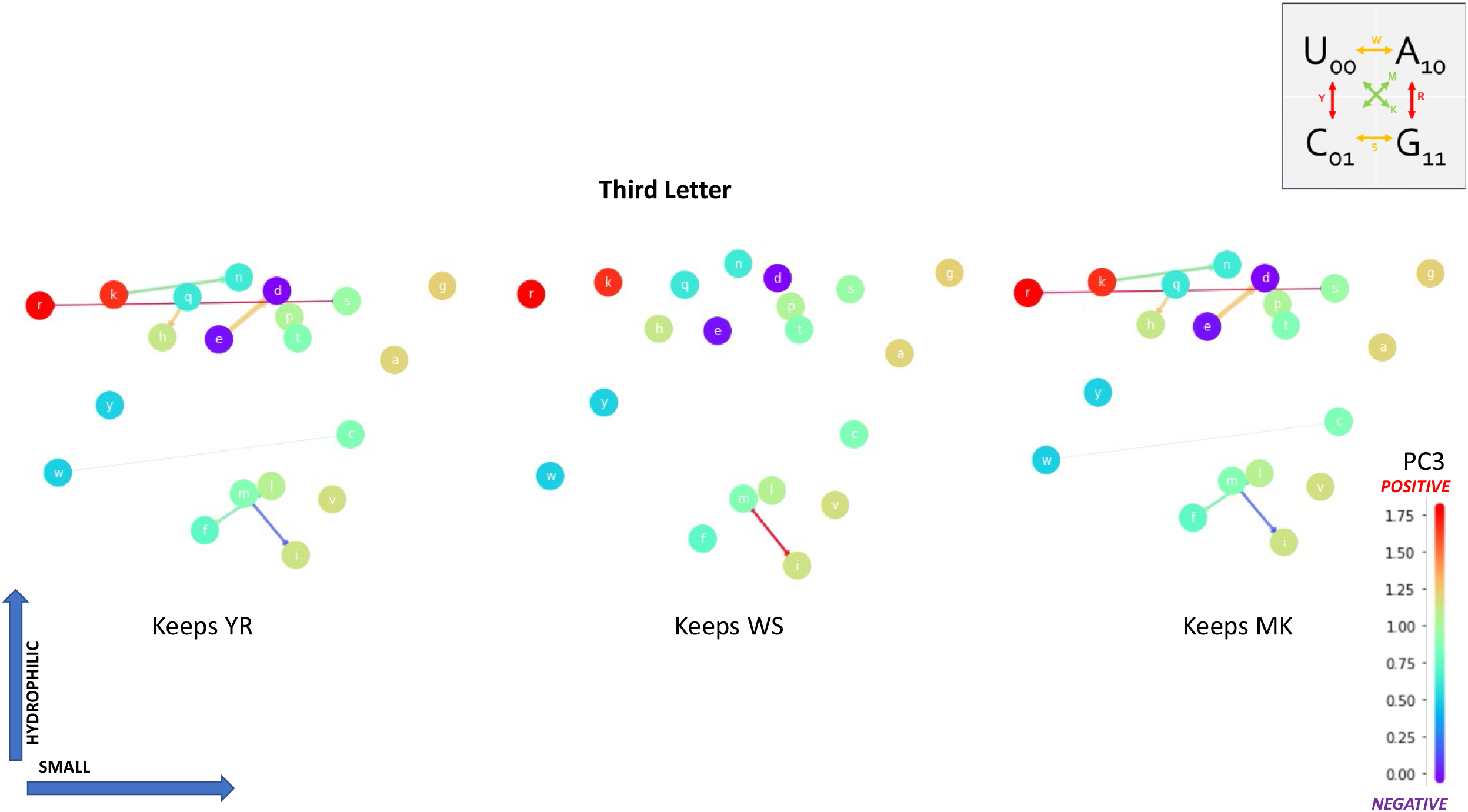
Substitutions sorted by mutation type. The change of base is classified by the property preserved. For example, a U <-> A mutation is Y because both are pyrimidines. The positions of the mutations are shown in decreasing order of importance: (a) middle, (b) first, and (c) last nucleotide of the codon. The color of the nodes shows the PC3 value of each amino acid. The width and color of the edges show the likelihood of occurrence and change of deleterious effect, respectively in a database of oncogenes.^58^

**Figure 5:**
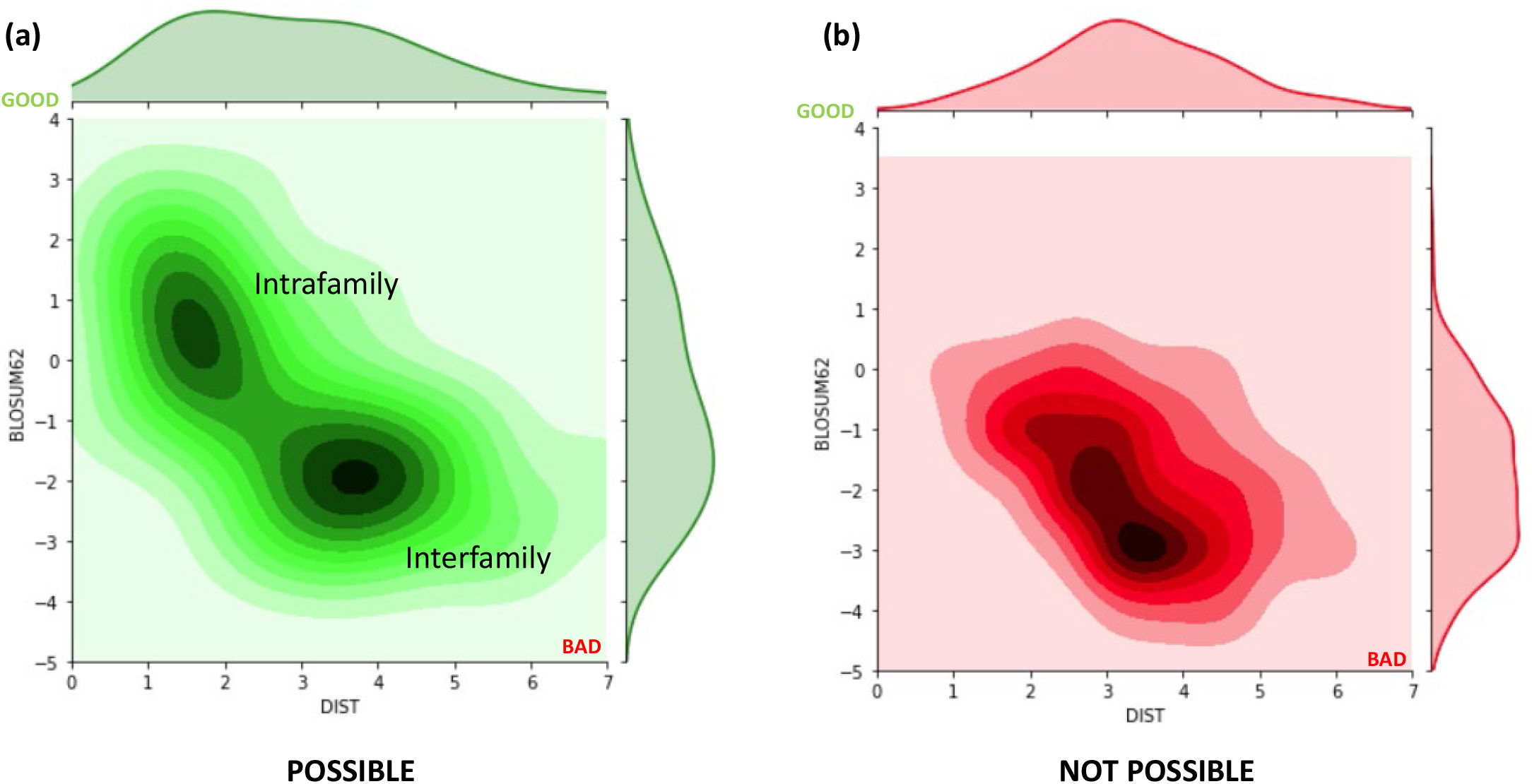
Heatmap of possible and not possible substitutions. The x-axis represents the length of the edge in the physicochemical space, while the y-axis shows the corresponding BLOSUM62 value, with negative values being more damaging.

**Table 1:**
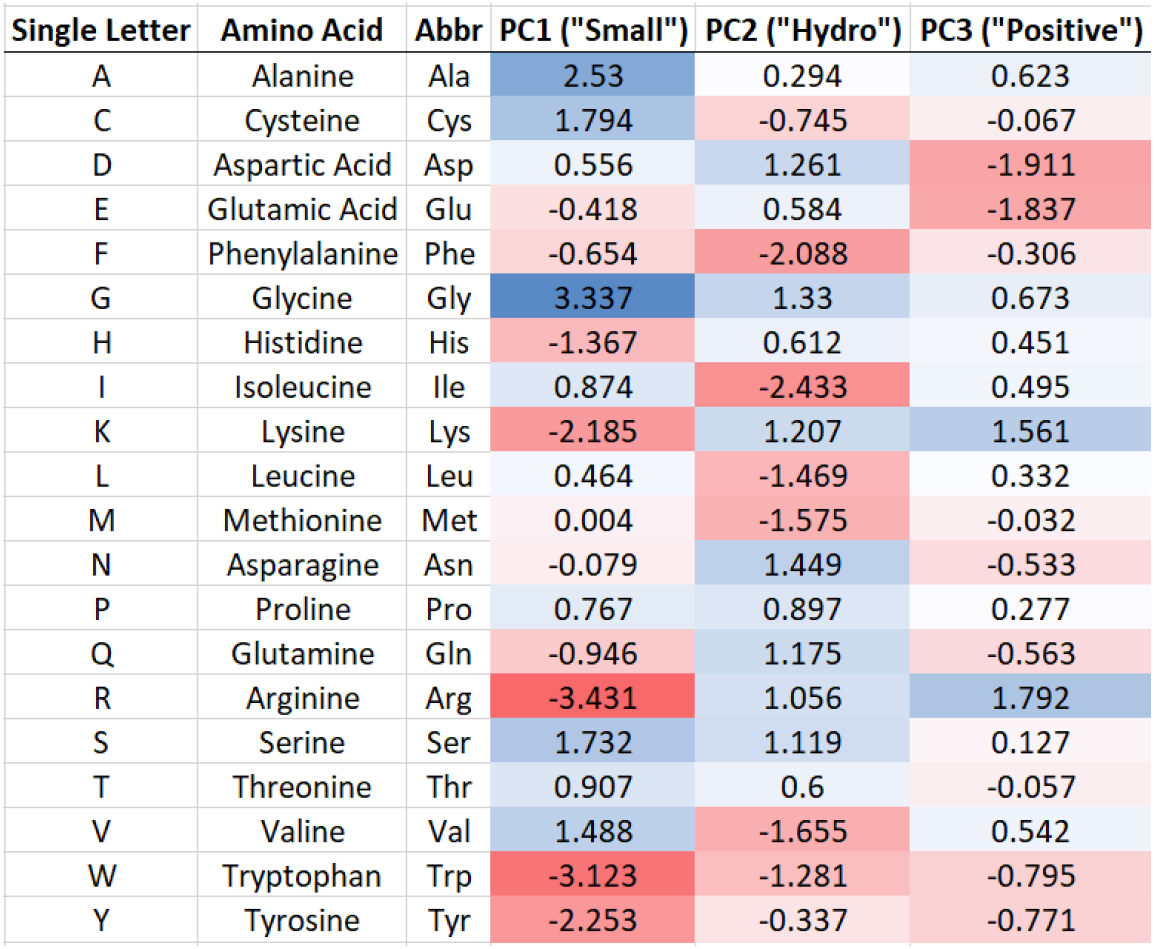
Amino acid names, abbreviations, and PCA values. For underlying data, see [ref 32]

**Table 2:** Number of mutations connecting each substitution pair.

|  | a | c | d | e | f | g | h | i | k | l | m | n | p | q | r | s | t | v | w | y |
| --- | --- | --- | --- | --- | --- | --- | --- | --- | --- | --- | --- | --- | --- | --- | --- | --- | --- | --- | --- | --- |
| a | 12 | 0 | 2 | 2 | 0 | 4 | 0 | 0 | 0 | 0 | 0 | 0 | 4 | 0 | 0 | 4 | 4 | 4 | 0 | 0 |
| c | 0 | 2 | 0 | 0 | 2 | 2 | 0 | 0 | 0 | 0 | 0 | 0 | 0 | 0 | 2 | 4 | 0 | 0 | 2 | 2 |
| d | 2 | 0 | 2 | 4 | 0 | 2 | 2 | 0 | 0 | 0 | 0 | 2 | 0 | 0 | 0 | 0 | 0 | 2 | 0 | 2 |
| e | 2 | 0 | 4 | 2 | 0 | 2 | 0 | 0 | 2 | 0 | 0 | 0 | 0 | 2 | 0 | 0 | 0 | 2 | 0 | 0 |
| f | 0 | 2 | 0 | 0 | 2 | 0 | 0 | 2 | 0 | 6 | 0 | 0 | 0 | 0 | 0 | 2 | 0 | 2 | 0 | 2 |
| g | 4 | 2 | 2 | 2 | 0 | 12 | 0 | 0 | 0 | 0 | 0 | 0 | 0 | 0 | 6 | 2 | 0 | 4 | 1 | 0 |
| h | 0 | 0 | 2 | 0 | 0 | 0 | 2 | 0 | 0 | 2 | 0 | 2 | 2 | 4 | 2 | 0 | 0 | 0 | 0 | 2 |
| i | 0 | 0 | 0 | 0 | 2 | 0 | 0 | 6 | 1 | 4 | 3 | 2 | 0 | 0 | 1 | 2 | 3 | 3 | 0 | 0 |
| k | 0 | 0 | 0 | 2 | 0 | 0 | 0 | 1 | 2 | 0 | 1 | 4 | 0 | 2 | 2 | 0 | 2 | 0 | 0 | 0 |
| l | 0 | 0 | 0 | 0 | 6 | 0 | 2 | 4 | 0 | 18 | 2 | 0 | 4 | 2 | 4 | 2 | 0 | 6 | 1 | 0 |
| m | 0 | 0 | 0 | 0 | 0 | 0 | 0 | 3 | 1 | 2 | 0 | 0 | 0 | 0 | 1 | 0 | 1 | 1 | 0 | 0 |
| n | 0 | 0 | 2 | 0 | 0 | 0 | 2 | 2 | 4 | 0 | 0 | 2 | 0 | 0 | 0 | 2 | 2 | 0 | 0 | 2 |
| p | 4 | 0 | 0 | 0 | 0 | 0 | 2 | 0 | 0 | 4 | 0 | 0 | 12 | 2 | 4 | 4 | 4 | 0 | 0 | 0 |
| q | 0 | 0 | 0 | 2 | 0 | 0 | 4 | 0 | 2 | 2 | 0 | 0 | 2 | 2 | 2 | 0 | 0 | 0 | 0 | 0 |
| r | 0 | 2 | 0 | 0 | 0 | 6 | 2 | 1 | 2 | 4 | 1 | 0 | 4 | 2 | 18 | 6 | 2 | 0 | 2 | 0 |
| s | 4 | 4 | 0 | 0 | 2 | 2 | 0 | 2 | 0 | 2 | 0 | 2 | 4 | 0 | 6 | 14 | 6 | 0 | 1 | 2 |
| t | 4 | 0 | 0 | 0 | 0 | 0 | 0 | 3 | 2 | 0 | 1 | 2 | 4 | 0 | 2 | 6 | 12 | 0 | 0 | 0 |
| v | 4 | 0 | 2 | 2 | 2 | 4 | 0 | 3 | 0 | 6 | 1 | 0 | 0 | 0 | 0 | 0 | 0 | 12 | 0 | 0 |
| w | 0 | 2 | 0 | 0 | 0 | 1 | 0 | 0 | 0 | 1 | 0 | 0 | 0 | 0 | 2 | 1 | 0 | 0 | 0 | 0 |
| y | 0 | 2 | 2 | 0 | 2 | 0 | 2 | 0 | 0 | 0 | 0 | 2 | 0 | 0 | 0 | 2 | 0 | 0 | 0 | 2 |

To perform a more quantitative analysis that captures the “directedness” of amino acid substitutions in the physicochemical space independent of distance, a measure of information entropy is used. The entropy will be zero if the change is entirely along one direction (PC1, PC2, or PC3), while it will be maximized if the change is equal in all three simultaneously. First, the components are normalized by the vector length squared:

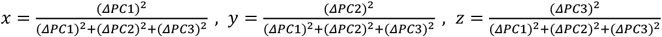

This ensures that x + y + z = 1. Then, the Shannon entropy^40^ is calculated:

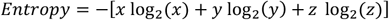

Finally, the entropy is divided by the maximum value,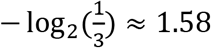, so the normalized entropy will always be between zero and one. Figure 6a separates the amino acid substitutions based on the number of codon pairs that differ by one nucleotide. For example, each of valine’s four codons (GUU, GUC, GUA, GUG) can be mutated to code for alanine (GCU, GCC, GCA, GCG) by changing the middle nucleotide from U to C. Therefore, the V <-> A substitution will be classified as having 4 codons. Substitutions that are not accessible have zero codons. The accessible changes are further separated into “low codon” for 1, 2, or 3 possible codons, and “high codon” for 4 or 6 codons. The normalized entropy distributions for the three different categories are shown in figure 6b. As a test case for using the tools introduced here, a dataset^41,42,43^ with 175,297 amino acid substitutions in the SARS-CoV-2 receptor binding domain is employed. For this visualization, the mean viral escape value for each substitution is calculated across all listed sites. Now, there are 337 datapoints, since forward and backwards substitutions are counted separately, but not every change appears in the dataset. The mean escape versus the normalized entropy or change in PC2 (the principal component that measures hydrophobicity) are plotted in figures 7a and 7b, respectively. Next, the physicochemical vectors representing substitutions associated with high and low mean escape are shown (figure 8). The edges are colored by the normalized entropy for each change. Finally, select results are visualized from another recent dataset^44^ by the same research group produced via site saturation mutagenesis experiments.^45^ Here, the change in binding of sarbecovirus spike proteins, including those of SARS-CoV-1 (SARS) and SARS-CoV-2 (COVID-19), with human ACE2 receptors are analyzed. In these assays, all possible amino acid substitutions at six analogous sites are measured.

**Figure 6:**
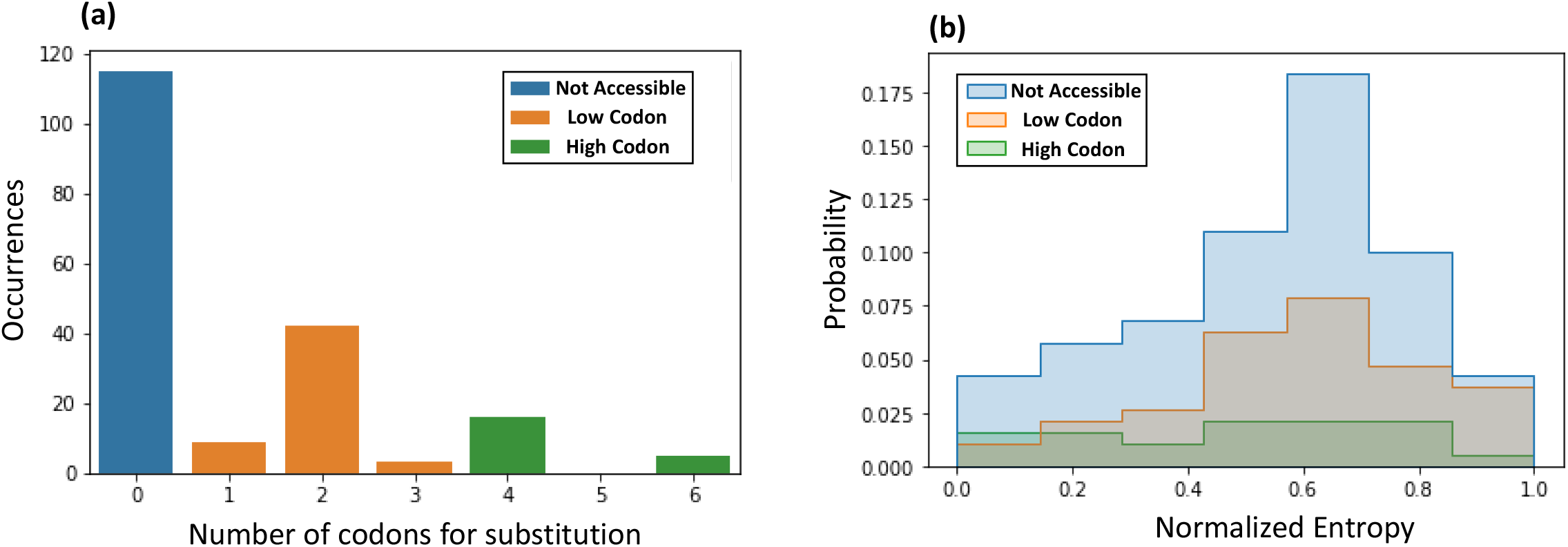
(**a**) Separation of amino acid substitutions based on the number of associated codons. “Not accessible” substitutions have zero codon pairs for the original and substituted amino acids that differ by a single nucleotide. If 1, 2, or 3 codon permutations exist, it is classified as “low codon,” and “high codon” if 4 or 6 exist. (**b**) The distributions of normalized entropy for each category of substitutions. Small values of normalized entropy correspond with substitutions that change just one direction at a time. On the other hand, large values show that multiple properties changed at the same time, which show up as diagonal lines in the physicochemical space visualizations.

**Figure 7:**
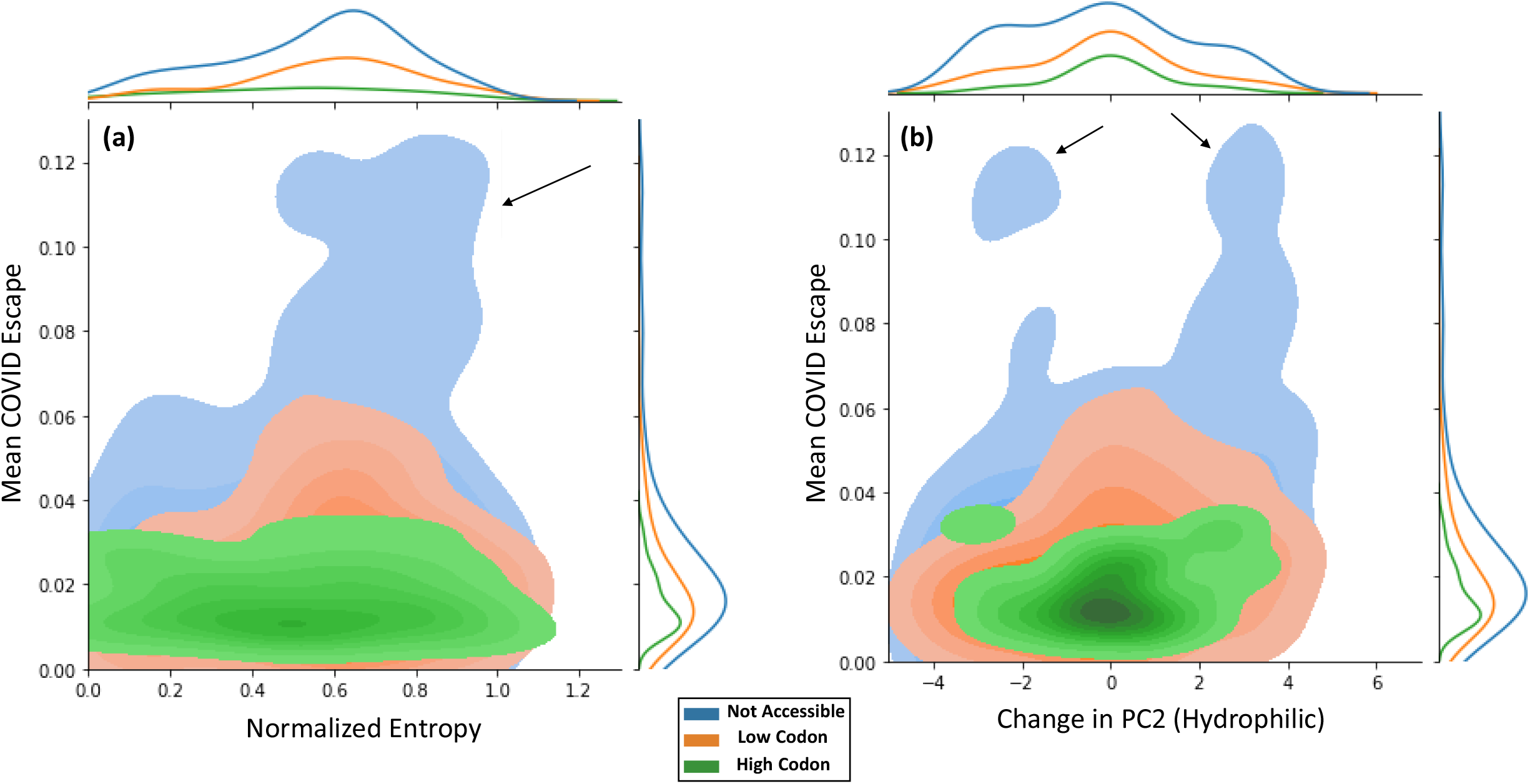
Mean escape values for SARS-CoV-2 receptor binding domain versus (**a**) normalized entropy of the substitution or (**b**) change in PC2, which represents the hydrophilic nature of the amino acids. The arrows, which indicate the substitutions associated with the largest immune escape, tend to have large normalized entropies and changes in PC2. They are also not accessible via single nucleotide mutations.

**Figure 8:**
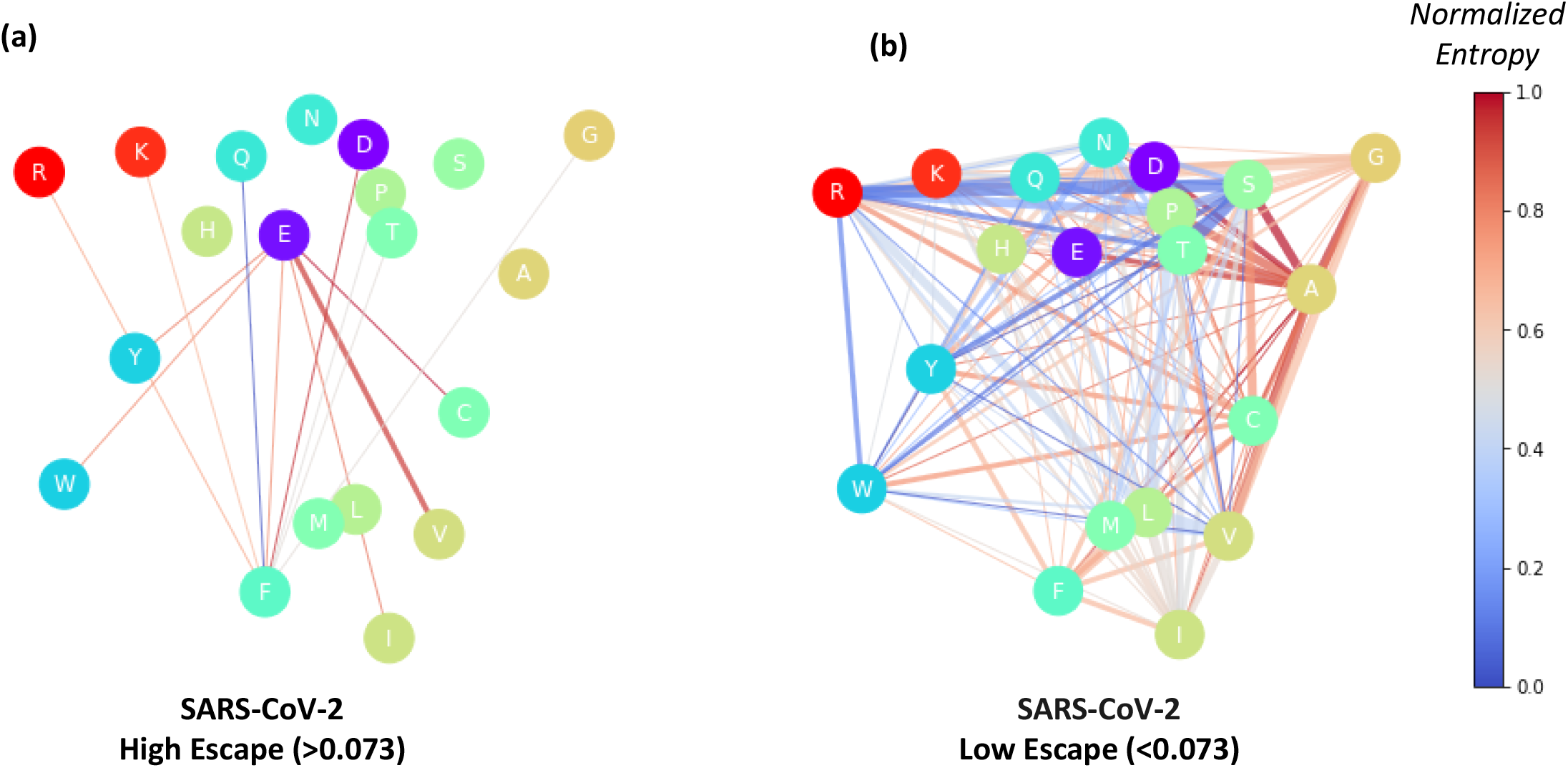
PCA visualizations showing the substitutions (**a**) above or (**b**) below the midpoint of the mean escape values in the dataset (0.073). The high-escape substitutions change multiple properties at once, as indicated by the diagonal lines and large normalized entropy values.

## Discussion of Results

The physicochemical vector space defined by the PCA shows a clear clustering of properties based on the middle nucleotide of the codon. This modularity is an initial hint that the genetic code should be viewed as a kind of choice architecture. Notably, if the middle letter is not changed by a mutation, it is likely that the substitution will be conservative, or even synonymous.

Figure 2 shows how the modularity extends to the accessible amino acid substitutions. There are many short connections between similar residues, and fewer long connections between amino acid families. This is even more noticeable when compared with the substitutions that are not possible via a single point mutation. There is also a significant difference in the orientation of the edges, in which possible substitutions correspond to lines more likely to horizontal or vertical, but not diagonal. This is associated with physicochemical changes confined to size or hydrophobicity, respectively, while a diagonally oriented edge changes both simultaneously. Because both positively (R, K) and negatively (E, D) charged residues are naturally hydrophilic and tend to be large, substitutions will be automatically constrained mostly within the z direction. As an illustration, K <-> E is possible and represents a very large change in charge, but has a BLOSSUM62 of +1, which indicates a benign change.

The single-nucleotide mutations shown in figure 3 help illustrate the effects of accessible substitutions. In some cases of heterozygous advantage, as with carriers of one sickle hemoglobin gene who are more resistant to malaria,^46^ a significant change in properties occurs even though it is caused by a point mutation. However, the change is limited in that sickle hemoglobin differs enough from the wild type to confer increase malaria resistance, but is similar enough so that symptoms only appear in homozygous individuals, and even then, only under low-oxygen conditions. In figure 4, the edges are separated by nucleotide position and mutation type. Here, mutations in the middle nucleotide (fig 4a) tend to be more vertically directed, while those of the first nucleotide are horizontally oriented (fig 4b). The sparsity of edges in fig 4c shows how few nonsynonymous mutations are possible by changing the third nucleotide. These findings lend themselves to the interpretation that the canonical genetic code is arranged such that the size and hydrophobicity of the residues can be changed mostly independently. Specifically, changing the middle nucleotide primarily impacts the hydrophobicity, while the size is controlled by the first letter of each codon.

Figure 5 looks at the Euclidean distance for possible and not possible substitutions. This approach is comparable to the Miyata^47^ or Grantham^48^ distance metrics. However, these measures use volume and polarity (along with composition in the Grantham case), which are correlated with each other, while here orthogonal principal components are employed. As expected, the distance and BLOSUM62 values are inversely related, showing that swapping very dissimilar amino acids is likely to be deleterious. Interestingly, the modular structure of the vector space enforced by the genetic code is apparent here too. The possible substitutions show distinct peaks for the short intrafamily and longer interfamily edges, while they are muddled into a single blob for the inaccessible substitutions. In figure 6, the impact of distance is removed, and instead, the directedness along one axis is quantified using the normalized entropy. The “high codon” substitutions differ in that a much larger fraction of them have low entropy, as compared with the inaccessible and “low codon” categories.

The substitutions showing the highest escape in figure 7a, as indicated by the arrows, all tend to have large normalized entropy values, which means they change multiple properties at once. In particular, they tend to have large changes in PC2, as seen in figure 7b. In the case of the dataset used, the highest escape substitutions involve glutamic acid (E) or phenylalanine (F). They also had high entropy, as indicated by the diagonal lines and colors. With phenylalanine, there may also the effect of altering the aromatic cage^49^ binding.

These results can be interpreted to indicate that the mutations that can best escape the immune system tend to change both the size and hydrophobicity at the same time. They also are not usually accessible by a single nucleotide mutation, as is the case with most large changes in physicochemical properties. Additionally, it may be adaptive for the host immune system to expend less resources on preventing escape in these situations, since the requirement that at least two mutations occur makes them inherently less likely.

Finally, the novel “fireworks” diagrams shown in figures 9 and 10 illustrate the usefulness of this approach to visualizing amino acid substitutions in physicochemical space. Binding with the ACE2 receptors in lung cells is often a crucial step in the infection cycle or respiratory viruses. Here, blue arrows represent strong binding, while red arrows represent impaired binding ability. The dashed lines show amino acid substitutions that are not accessible by a single nucleotide mutation, and the thickness of each solid line indicates the number of associated codons. Even though the pathogens responsible for SARS (SARS-CoV-1) and COVID-19 (SARS-CoV-2) are closely related, it is known that mutations at analogous sites can have very different effects on ACE2 binding affinity. As seen in figure 9, mutations at position 501 that produce a positively charged amino acid (R or K) significantly reduce binding between the SARS virus and ACE2, as will a mutation to Y. In contrast, the binding of some SARS-CoV-2 variants is greatly enhanced by a N501Y substitution.^50^ In both cases, the downward blue lines indicate that substitutions to more hydrophobic amino acids do not inhibit binding (except for L).

**Figure 9:**
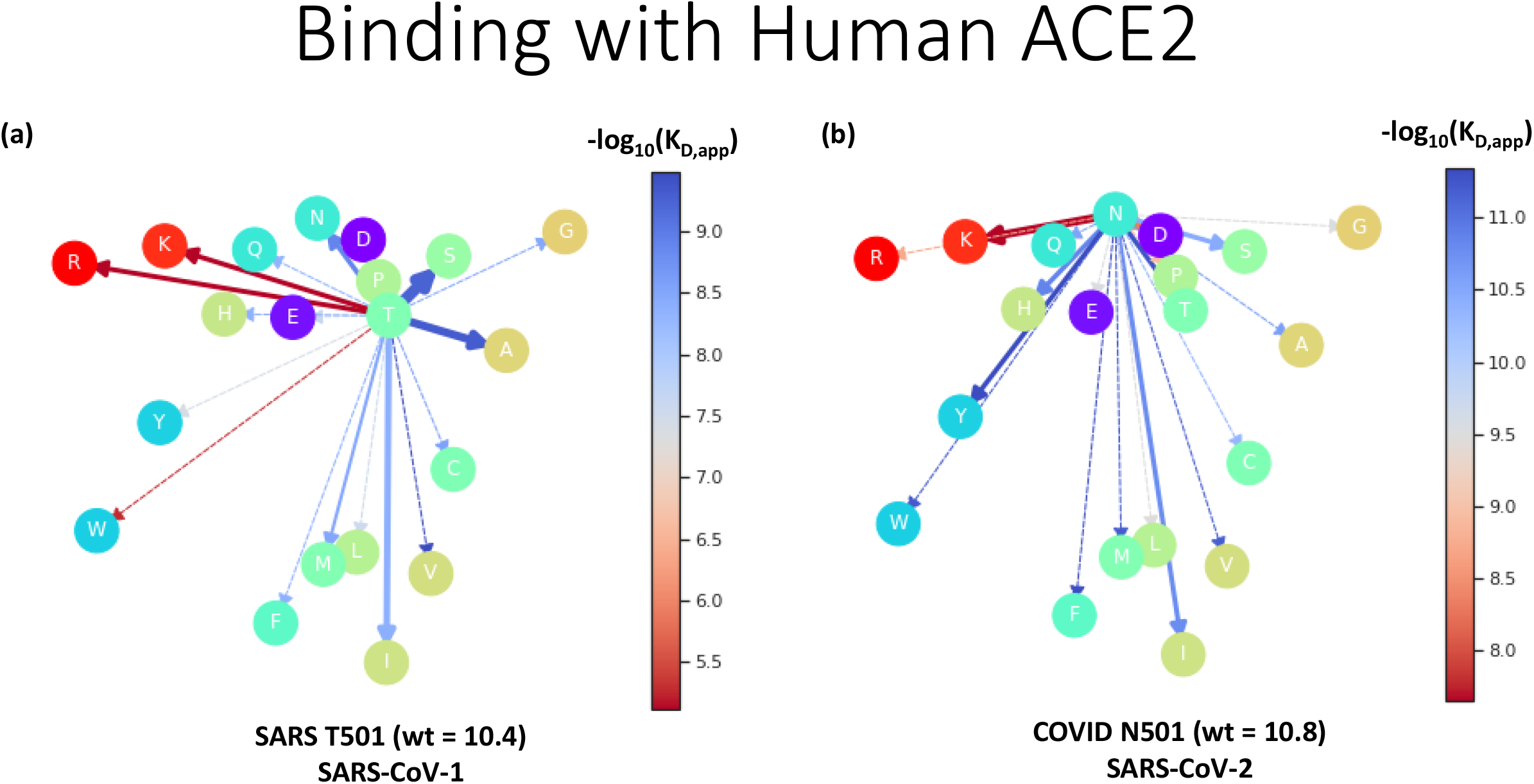
“Fireworks graphs” from site saturation mutagenesis experimental data. Blue and red lines show high and low binding affinity with human ACE2, respectively. Thicker lines indicate more associated codon permutations, and dashed lines show substitutions that are not accessible by single nucleotide mutations. Here, a (**a**)SARS-CoV-1 (SARS) and a (**b**)SARS-CoV-2 (COVID-19) variant at site 501 are shown. In the case of SARS, substitutions to large, and especially positively charged, amino acids disrupt binding the most. For COVID-19, the most damaging changes are to the positively charged R or K, and only the latter is accessible via point mutation.

**Figure 10:**
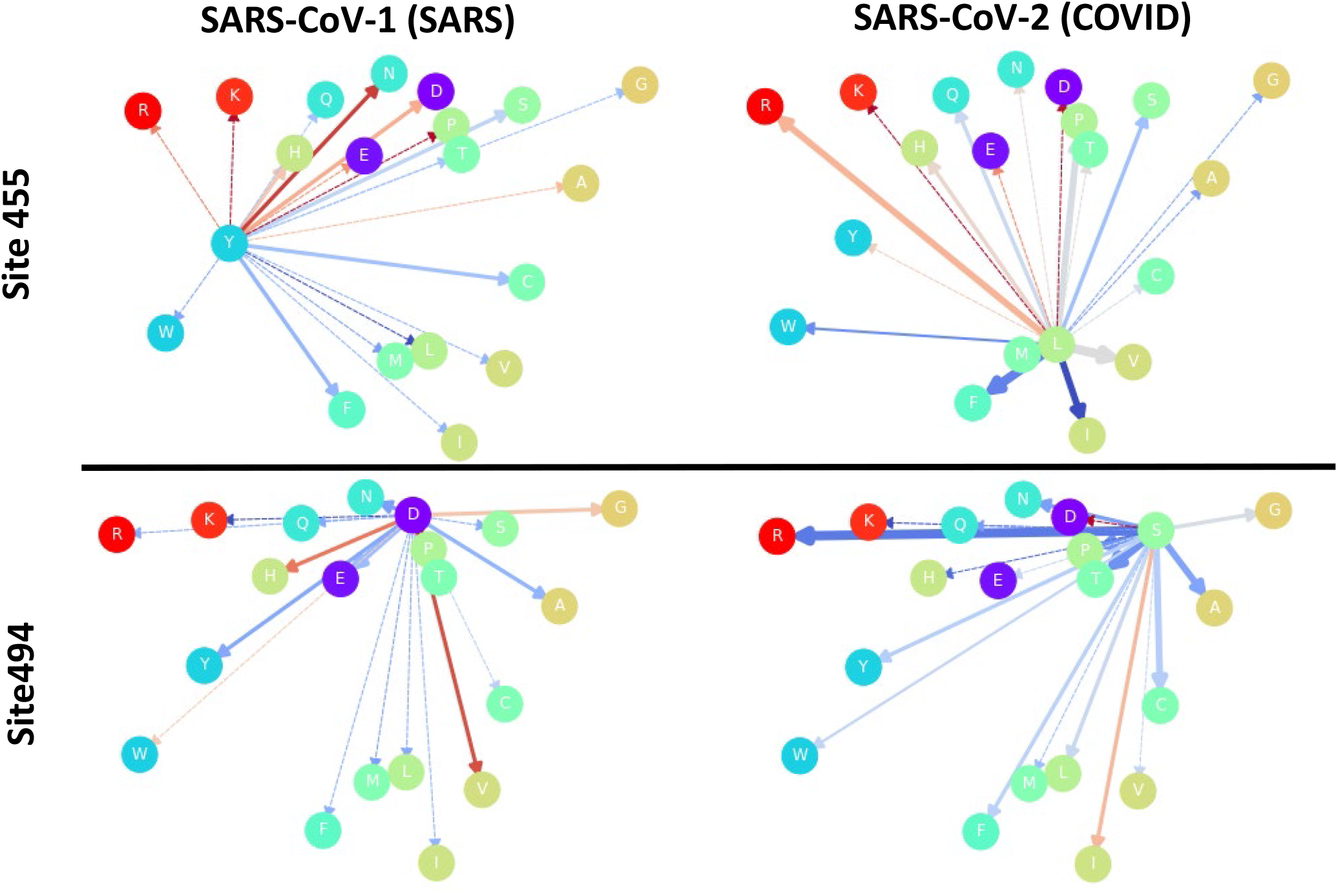
Fireworks graphs for SARS-CoV-1 and SARS-CoV-2 variants at sites 455 and 494. Sometimes, binding inhibition is similar across sectors of related amino acids, as for the large hydrophilic residues at site 455. In other situations, the changes are limited to particular substitutions, as seen for the data for site 494.

To further highlight some comparisons of analogous sites in the dataset, figure 10 places sites 455 and 494 side-by-side for SARS-CoV-1 and SARS-CoV-2. It can be seen that the SARS virus is susceptible to reduced binding if Y455 is replaced by any of a cluster of amino acids that are large and hydrophilic. This not that case for the COVID-19 virus at the analogous site (L455) since it will suffer significantly reduced binding only if replaced with a charged amino acid. Interestingly, many of these changes are precluded from occurring by a single point mutation. At site 494, both SARS and COVID-19 viral binding apparently suffer only due to specific substitutions, as opposed to sectors of amino acids with similar properties. For example, D494V is accessible by a single-nucleotide mutation, and has a large negative impact on the binding between SARS-CoV-1 and ACE2. A similar particularized substitutions for SARS-CoV-2 would be S494I.

The current work focuses specifically on missense point mutations allowed by the canonical genetic code to elucidate this particular set of explicit constraints. Other types of mutations, such as nonsense mutations that lead to premature termination of the protein, as well as base deletions or insertions, are not considered here, since they depend on many contextual factors, such as adjacent codons. Similarly, there is increasing evidence that even synonymous mutations can create variation subject to selection via differences in gene expression and protein folding.^51^ It has also been proposed that “hidden stops”^52^ serve as an additional fail-safe mechanism to halt the translation of faulty proteins in the event of mutation. These would be another example of a choice constraint intrinsic to the canonical genetic code, but are beyond the scope of the current work. Nonstandard codes are minor modifications stemming from standard code, and often involve assigning an amino acid to a stop codon.^53^

## Conclusions

The canonical genetic code places constraints on the amino acids substitutions that are accessible via single-nucleotide point mutations. Just as transcribing text with a typewriter^54^ is more likely to lead to usable variation compared with either a perfect fidelity copy-machine or a leaky pen, the choice architecture encountered by random mutations is crucial to the generation of bounded adaptation. Here, it is proposed that in addition to simple error minimization, the genetic code contributes to the evolvability of new variation by permitting mostly independent control over the size and hydrophobicity of amino acids, which are the two most important physicochemical properties. This can have clinical applications for heritable disorders, as well as implications for evolutionary theory. In the current era of large genomic data, the use of machine learning, including the identification of heritable conditions due to single-nucleotide polymorphisms, can be informed by the findings of this work. The constraint that mutational trajectories^55^ consist only of changes accessible via point mutations may make the evolution of certain traits more predictable.^56^ Also, the rational design of new proteins and vaccines can potentially be accomplished more efficiently, leading to new therapeutic interventions. In particular, the immune escape of viral variants with single or multiple mutations can be more easily predicted and prevented. Finally, new methods of visualizing and presenting data, as with the novel “fireworks” diagrams, can help make sense of large datasets, such as those produced by site saturation mutagenesis or other modern bioinformatics techniques.

## Acknowledgements

I would like to express my sincere appreciation to Rees Kassen for many enlightening discussions. This work was supported by NSU PFRDG #335472.

## Notes

https://github.com/jbloomlab/SARSr-CoV_homolog_survey/blob/master/results/final_variant_scores/mut_variant_scores.csv

### Competing Interest Statement

The authors have declared no competing interest.

### Summary of Updates

Revised August 2022

